# CLARUS: An Interactive Explainable AI Platform for Manual Counterfactuals in Graph Neural Networks

**DOI:** 10.1101/2022.11.21.517358

**Authors:** Jacqueline Beinecke, Anna Saranti, Alessa Angerschmid, Bastian Pfeifer, Vanessa Klemt, Andreas Holzinger, Anne-Christin Hauschild

## Abstract

**Background:** Lack of trust in artificial intelligence (AI) models in medicine is still the key blockage for the use of AI in clinical decision support systems (CDSS). Although AI models are already performing excellently in systems medicine, their black-box nature entails that patient-specific decisions are incomprehensible for the physician. This is especially true for very complex models such as graph neural networks (GNNs), a common state-of-the-art approach to model biological networks such as protein-protein-interaction graphs (PPIs) to predict clinical outcomes. The aim of explainable AI (XAI) algorithms is to “explain” to a human domain expert, which input features, such as genes, influenced a specific recommendation. However, in the clinical domain, it is essential that these explanations lead to some degree of causal understanding by a clinician in the context of a specific application.

**Results:** We developed the CLARUS platform, aiming to promote human understanding of GNN predictions by allowing the domain expert to validate and improve the decision-making process. CLARUS enables the visualisation of the patient-specific biological networks used to train and test the GNN model, where nodes and edges correspond to gene products and their interactions, for instance. XAI methods, such as GNNExplainer, compute relevance values for genes and interactions. The CLARUS graph visualisation highlights gene and interaction relevances by color intensity and line thickness, respectively. This enables domain experts to gain deeper insights into the biological network by identifying the most influential sub-graphs and molecular pathways crucial for the decision-making process. More importantly, the expert can interactively alter the patient-specific PPI network based on the acquired understanding and initiate re-prediction or retraining. This interactivity allows to ask manual counterfactual questions and analyse the resulting effects on the GNN prediction.

**Conclusion:** To the best of our knowledge, we present the first interactive XAI platform prototype, CLARUS, that allows not only the evaluation of specific human counterfactual questions based on user-defined alterations of patient PPI networks and a re-prediction of the clinical outcome but also a retraining of the entire GNN after changing the underlying graph structures. The platform is currently hosted by the GWDG on https://rshiny.gwdg.de/apps/clarus/.

## 1 Introduction

Machine learning (ML) and artificial intelligence (AI) have become part of many areas of our everyday life. However, the transfer of such technologies to the medical domain, in particular in the form of clinical decision support systems (CDSS), is often presented with significant challenges. In domains where decisions have the potential to impact individuals’ health, there is an increased necessity to establish trust in employed AI models. The application of AI in the medical domain is particularly critical since it often involves patient care or treatment development [33]. The lack of interpretability in AI within such a medical settings is thus considered unacceptable due to the risk of erroneous predictions leading to severe consequences [8]. In particular, the blackbox nature of complex AI models often contributes to this lack of interpretability and, consequently, undermines trust in AI systems as CDSS[35].

In recent years explainable AI (XAI) models have been developed to help build trust in AI models and increase their transparency [16]. They achieve this, for example, by calculating relevance scores for inputs based on the AI models configurations [43][2], by creating saliency maps [40] or by analysing the model down to a single neuron [9]. Several surveys and practical tutorials compare and analyse state-of-the-art XAI models in different scenarios with different types of data [3] [19].

In the medical and biomedical domain, however, transparency alone is not sufficient in reaching a state of “explainable precision medicine”. In medicine, there is a need to move from explainability to causability [17], the measurable extent to which an explanation from an XAI model achieved a specified level of causal understanding. In the biomedical domain, this entails, for instance, the need to transition from exclusively focusing on gene importance to a causal understanding of the effects incurred by the absence or alteration of a gene.

In order to connect the transparency of XAI models with the need for causability, we need interactive XAI platforms that guide domain experts through the explanations given by an XAI model. Such platforms will allow the expert to gain insight into what influenced the AI model during the decision-making process. Additionally, it can help to identify confounding factors that might inhibit model performance or correlated factors that are biologically meaningful or inconsequential. Most importantly, it is imperative to allow the expert to interactively ask counterfactual questions (“if-not”, “why-not”, and “what-if” questions) and change their data based on those. Ultimately, this interaction will help to see how changes affect not only the AI model but also the XAI model, hence, increasing their causal understanding. There is a plethora of research works that deal with the automatic generation of counterfactual explanations for graphs [30]. Most of them try to generate the counterfactuals automatically, for example with the use of generative models [22], by just concentrating on edge deletion [21] or by a search driven by a Reinforcement Learning (RL) algorithm [26]. The main goal of those methods is to form a provided graph instance to compute another that has a small graph edit distance and sparsity (modification of fewer features as possible) in the smallest possible time. Preliminary research tries to compare several graph counterfactual generation frameworks under many datasets and evaluation metrics [31], and the result shows no clear winner in all metrics. The methods listed and evaluated do not incorporate solutions where the graph counterfactual generation was driven by human decisions. The evaluation of human-in-the-loop search strategies by similar means as the automated ones is one of the long-term goals of our research.

In most research domains, there are strong dependencies between entities such as genes that further impact their relation to the subsequent target parameters, like a clinical category. Such dependencies are often represented in graph structures. For example, in the medical and biomedical domain, much research is being done on protein-protein interaction (PPI) networks, especially in cancer research and subtyping, since gene mutations and other genetic alterations can lead to a gain or loss of PPIs [13]. Graph neural networks (GNN) have been specifically developed to handle graph-structured input data [50]. They have already been used on PPI networks, for example, to predict novel cancer genes [38] or for disease subnetwork detection [29]. Many XAI models for GNNs have already been developed, such as the GNNExplainer [53], PGMExplainer [47], XGNN [54], GNN-LRP [37], and GLRP [6]. But, as pointed out earlier, while these XAI models might increase the transparency of GNNs and help with understanding their inner processes, there is still a lack of causability.

For this reason, we developed CLARUS, an interactive XAI platform for GNNs. CLARUS will allow experts in the biomedical domain to understand why GNNs reach a particular decision and what features (genes) or interactions are important in the decision-making process. The prototype allows to interactively investigate three example PPI networks, namely, a synthetic network and two realworld networks for classifying Kidney renal clear cell carcinoma (KIRC). The interactivity will enable the expert to act based on their already gained understanding. Furthermore, CLARUS will illustrate the consequences of the action, and thus increase their causal understanding. CLARUS will pave the way for informed biomedical decision-making and the application of AI models in combination with various Omics datasets as CDSS.

## 2 Implementation

### 2.1 CLARUS framework

CLARUS is build with a fronted, backend, and an API for communication between the two. We developed the UI using R version 4.1.0 [32] and R Shiny [34] with important packages such as visNetwork [5] and igraph [7] for the graph visualisation and jsonlite [27] and httr [48] for the communication via an API. Screenshots of the UI can be found in the supplementary. The backend and API are implemented using Python 3.8 [46] using packages such as PyTorch [28] and PyTorch Geometric [10] for working with GNNs and Flask [11] for the API. Figure 1 shows an overview of what the user can expect from CLARUS.

**Figure 1:**
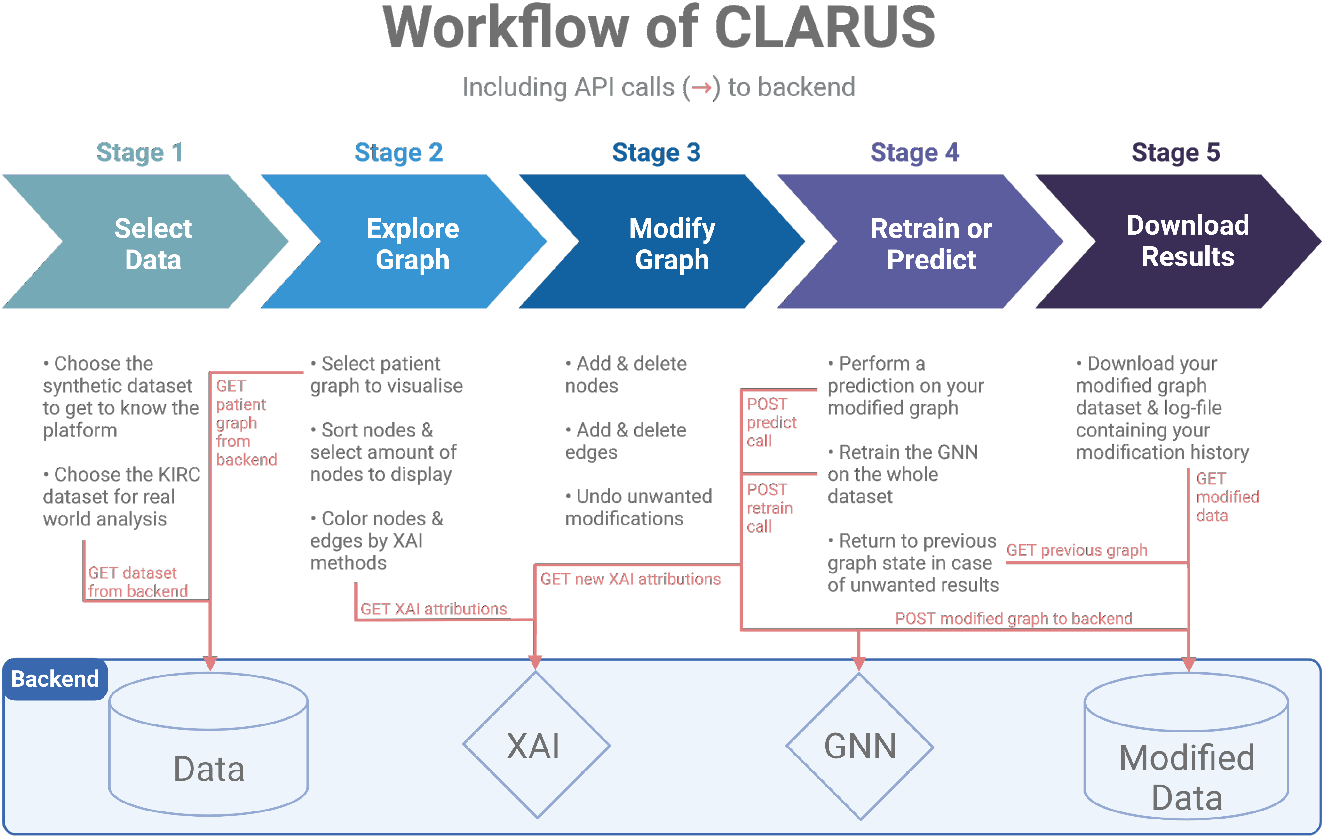
User workflow of CLARUS platform including API calls (red arrows) to the backend. The stages represent the main actions the user can perform on the UI. Namely, selecting one of the two datasets provided, exploring and modifying the visualised graph, performing a retraining or new prediction on the provided GNN, and lastly downloading the results. Stages two to four can be repeated until the user arrives at a desired result.

CLARUS provides a synthetic dataset which can be used to get to know the platform and its features, as well as, a real world PPI dataset on kidney cancer.

After selecting a dataset, the user can explore the visualised data alongside the performance of a pre-trained GNN. XAI methods such as GNNExplainer [53] can be selected at this stage to colour the nodes and edges of the data, giving the user deeper insights into the relevant features in the GNNs decision making process.

Based on their gained knowledge the user can then ask counterfactual questions and then manually manipulate the graph. After changing the graph, a new prediction on this graph can be initiated, or even a complete retraining of the GNN.

The user can then investigate the effect of their changes on the GNN prediction and XAI relevances. This will allow them to gain much deeper insights into the relevant features and the GNNs decision making process.

CLARUS is being hosted by the GWDG and reachable under the following domain: http://rshiny.gwdg.de/apps/clarus/

### 2.2 Dataset generation and preprocessing

#### 2.2.1 Synthetic dataset

To build the synthetic graph dataset we have generated 1000 Barabasi networks comprising 30 nodes and 29 edges. The networks had the same topology, but with varying node feature values. The features of the nodes were randomly sampled from a normal distribution *N*(0, 0.1). For one half of the networks we have selected two connected nodes to which we assigned values from *N*(1, 0.1). The GNN classifier identified the two specified classes with high accuracy. However, the ultimate goal here was to detect the selected nodes as the discriminatory factors Explainable AI methods should uncover these patterns in an algorithmic way. We investigate to what extend the human-in-the-loop could improve this process.

#### 2.2.2 KIRC dataset

To illustrate the usability of CLARUS in the biomedical domain we utilize an aggregated gene expression and methylation data set of cancer patients, as presented in [29]. The original datasets were downloaded from The Cancer Genome Atlas (TCGA) database^1^, which represents one of the largest collections of multi-omics data sets. It contains molecular and genetic profiles for over 33 different cancer types from 20,000 individual tumour samples [41].

The acquired data sets were harmonised so that for every patient multi-omics information was available. Furthermore, we focused on cancer-relevant genes as proposed by [39]. Genes containing missing values at least for one patient were removed from the analyses. The obtained numerical data matrices containing expression and methylation were normalised using *min-max* normalisation.

For our CLARUS example application we focus on a binary classification task differentiating multi-modal molecular footprints of patients with Kidney renal clear cell carcinoma (KIRC) and a random set of patients suffering from multiple types of cancer. As presented in Pfeifer et al. in 2022, each patient is represented by a patient-specific PPI network, where nodes or genes are enriched by feature values from DNA methylation and gene expression. The interaction information for the PPI network was retrieved from the STRING database [45].

In order to provide a smaller overview dataset, we employed the disease subnetwork discovery approach as presented in [29], to extract a cancer-specific disease module. We utalize this dataset, to demonstrate the different functionalities of the CLARUS prototype as described in section.

### 2.3 Model training and evaluation

Graph Convolutional Networks (GCNs) have been specifically introduced to handle graph type data such as PPI networks. They operate in a comparable manner to Convolutional Neural Networks, by aggregation of information from a neighbourhood of nodes and edges instead of a grid of pixels [20]. Through this aggregation, they are able to perform tasks such as link prediction, node classification, and graph classification. Since the training of GCNs on our datasets can take a few minutes, we have saved pre-trained GCNs for our datasets, which will be loaded upon selection of a dataset. This way, we have reduced the start-up time of CLARUS drastically

For each dataset, a random split into 70% training and 30% validation data has been performed. This split is fixed by a seed, such that the same graphs will always be used for retraining. Early stopping has been integrated into the training to avoid over-fitting. Prediction confidences for each graph, as well as performance of the GCN on the validation data in terms of sensitivity, specificity, and confusion matrix have been saved for the pre-trained models.

Buttons for prediction and retraining have been implemented (see Figure 2). A prediction will only prompt an update of the predicted label and the prediction confidence of the GCN, as well as, a recalculation of the XAI attributions. This takes only a few seconds. Retraining on the other hand might take a bit longer and will additionally lead to an update in the confusion matrix, sensitivity and specificity of the GCN.

**Figure 2:**
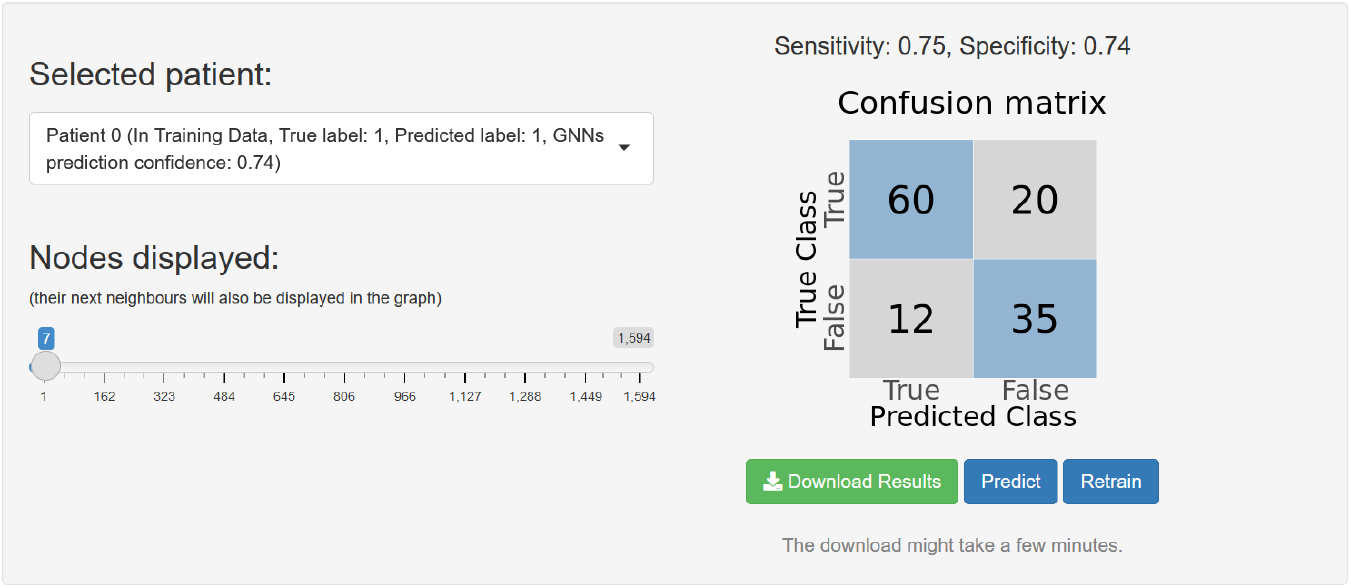
UI element of CLARUS showing (1) drop-down field for selecting patients (contains patient label number, true label of the patient, predicted label by the GCN, and prediction confidence of the GCN), (2) sliding bar for selecting how many nodes to display in the graph visualisation (see Figure 6), (3) confusion matrix alongside sensitivity and specificity values, and (4) buttons for predicting, retraining or downloading of results.

### 2.4 Visualisation of PPI networks with XAI attributions for genes and PPIs

To help the user gain insights into the molecular mechanisms within a patient graph and identify important genes or sub-networks, we have developed a graph visualisation that highlights node and edge relevances from a multitude of XAI models. For simplicity, we will use the terms nodes and edges as synonyms for genes and PPIs.

Two attribution methods were used to calculate positive attribution values on the edges [44] provided by the Captum library of Pytorch^2^. The saliency method computes for each edge the absolute value of the gradient of the non-linear function that the GNN computes w.r.t. the weight of this edge (as defined by the training process). The Integrated Gradients (IG) method is a more sophisticated version of the saliency method, where instead of the absolute value, the integral of the gradient w.r.t a range of learned weight values is used. The GNNExplainer [53] computes the important subgraph *GS* of the computation graph *Gc* [12], [49], [20] of the input graph *G* with the use of an optimisation algorithm. The algorithm is driven by the maximisation of the mutual information [24] between the tried substructure (*GS* of *Gc*) w.r.t. the predicted label distribution. The user can choose which XAI method to use for colouring the edges and nodes, representing the genes and PPIs respectively. The edges will be coloured in grey-scale based on the relevance values, while the nodes will be coloured in a blue-scale. The identification of important genes in the complete PPI network is often unfeasible if the number of genes is very large. Thus we have implemented a sliding bar (see Figure 2) that allows the user to select the top genes according to a specified measure, such as relevance values, degree or alphabetically. This ensures that the user can analyse the most important genes or if desired the most irrelevant genes. Alongside those geness their next neighbours (direct interactions partners) will also be displayed, to allow the user to see the interactions of the selected genes.

Furthermore, we implemented a tooltip that gets shown if the curser hovers over an edge or node. For edges, the tooltip will display information on which nodes the edge connects, as well as all relevances for this edge, labeled by the XAI model that computed them (see Figure 3). For nodes, the tooltip displays the node label and degree alongside all relevances for this node, labeled by the XAI model that computed them (see Figure 4).

**Figure 3:**
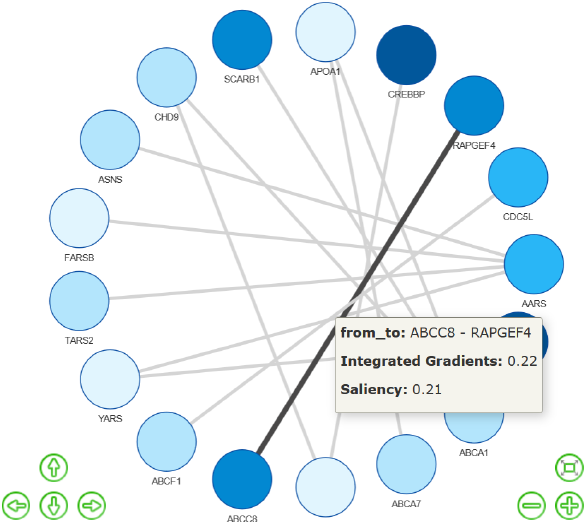
Edge tooltip: Hovering over an edge shows information on connected nodes and relevance values from XAI models, namely Integrated Gradients and Saliency methods.

**Figure 4:**
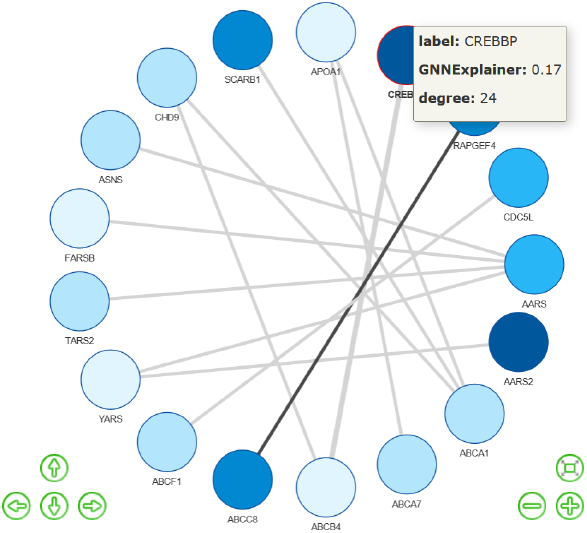
Node tooltip: Hovering over a node shows information on label, degree and relevance values from the GNNExplainer method.

In addition to the information in the tooltips, the user can find more details and features of the selected nodes and displayed edges in two data tables (see Figure 5).

**Figure 5:**
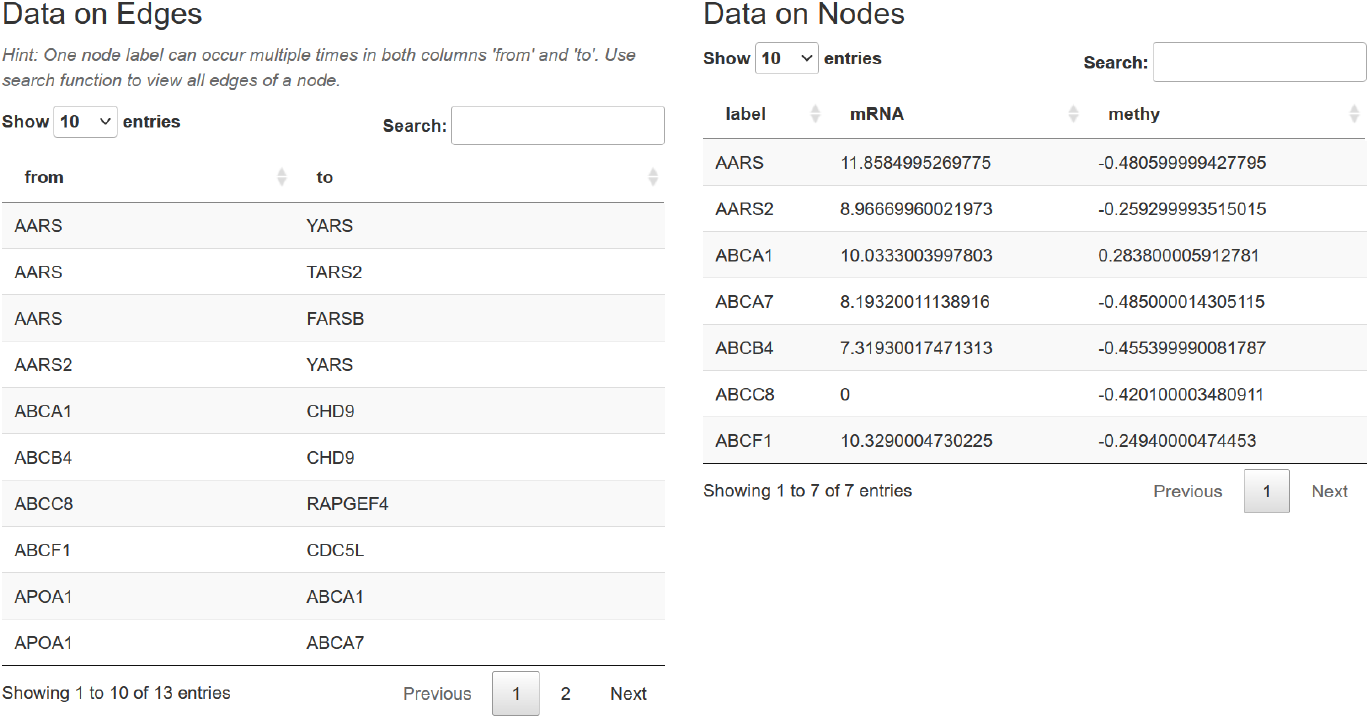
UI feature showing information on nodes and edges in the selected dataset.

### 2.5 Data manipulation

Besides network visualisation, one of our main contributions is the interactive manipulation of the of the patient-specific PPI networks and the demonstration of the impact of such manipulations. It allows the user to directly ask manual counterfactual questions by changing the network and answer them directly by updating the prediction or retraining the GNN and observing the effects of the counterfactual in the GNN performance and the newly calculated XAI relevances.

In order to allow counterfactual questions such as “What would have been the prediction if this node or edge would not have been in the network?” or “What would have been the model performance if this node or edge would have been present in the network?”, we have implemented the deletion and addition of nodes and edges. The user can manipulate the network by adding or deleting a multitude of nodes and edges and subsequently start retraining or updating the prediction to evaluate whether the changes resulted in an expected outcome. An undo function allows reversing all manipulations. After predicting or retraining, the current state of the network and the GNN will always be saved, allowing the user to jump between saved states later on. This ensures that the user can always return to a state before a prediction or retraining if the changes are not effective or beneficial. Figure 6 shows the network visualisation and manipulation features in the UI.

**Figure 6:**
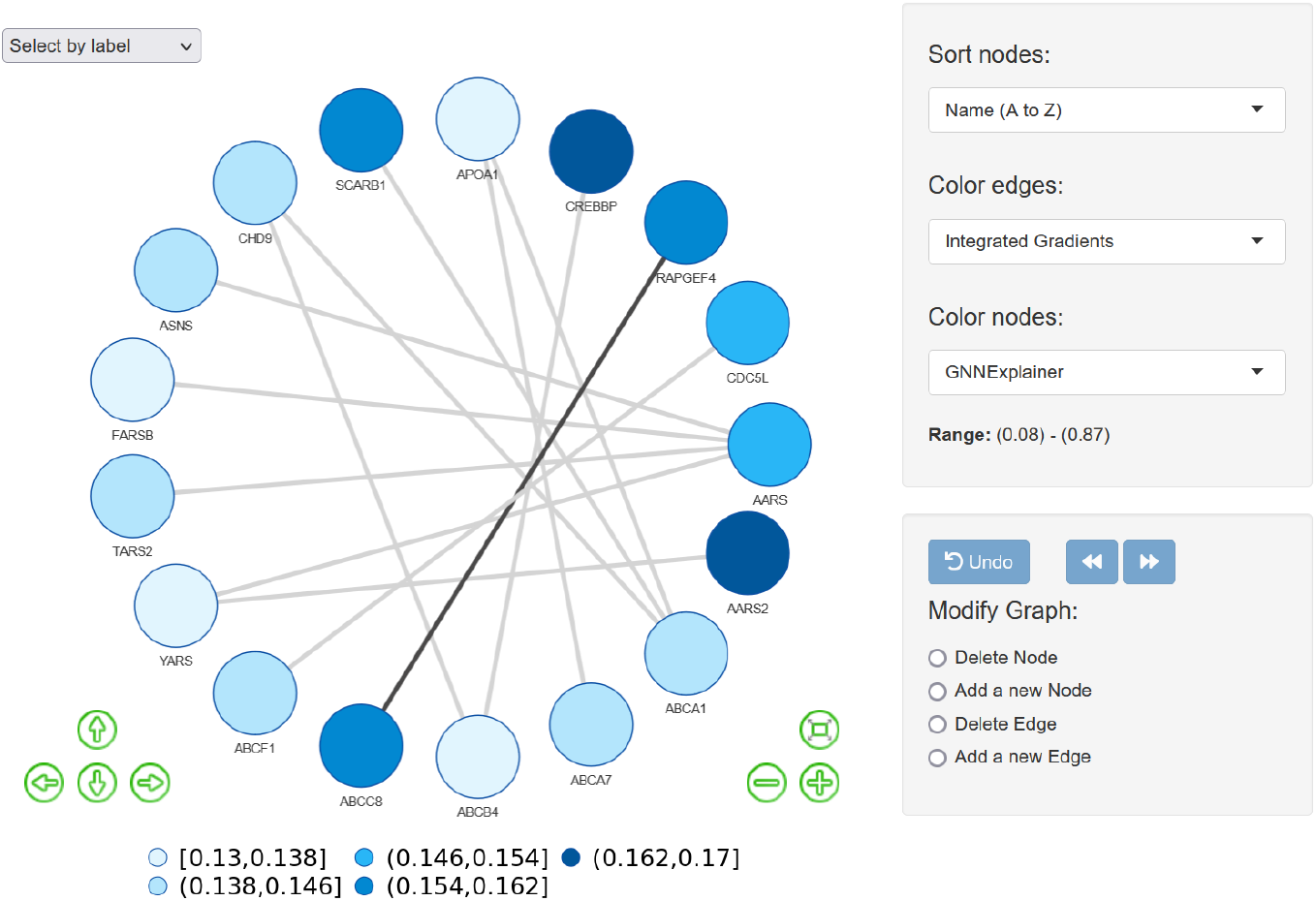
UI features of network visualisation and network manipulation. Left side: The visualised network with nodes coloured by GNNExplainer relevances, edges coloured by Integrated Gradients relevances, and a colour legend for node colours. Right side: Drop down menu’s that allow selecting on how to sort nodes, what values to use for colouring nodes and edges, as well as, information on the range of node relevance values. Beneath that are the radio buttons that enable addition and deletion of nodes and edges. The undo button is enabled as soon as the user performs one of the 4 manipulations and then allows to undo them. The forward and backward buttons are enabled when the user created a new network state, by performing a retraining or prediction on a manipulated network, and they allow the user to then move between the saved network states.

To demonstrate the effects the changes have on the GNN performance the user is shown a confusion matrix along with sensitivity and specificity of the GNN on the test data. The evaluation allows for a direct comparison of values such as true positives, false positives, true negatives and false negatives, while sensitivity and specificity identify the proportion of correctly identified positive and negative classes (see Figure 2). This enables a fast inspection to what degree the model performance was effected by the manual changes. The GNN performance of the initial training and every retraining will be printed in the log-file, allowing for an easy comparison.

In addition, the user can observe the effects of their changes by identifying if the predicted label and the confidence in the predicted label have changed.

Lastly, after retraining or predicting with the changed network the user will be able to identify changes in the relevance values. This shows that CLARUS not only enables the user to understand what consequences his changes have on the GNN performance, but also to understand the effects on the decision-making process of the GNN. By viewing changes in the relevances of nodes and edges, the user can gain a deeper knowledge on the correlations of the nodes and edges in the data and their influence in the decision-making process of the GNN. This should result in an increase in causability [25] and in new AI-interfaces [18], but of course this will need to be measured in future research.

### 2.6 Testing

Special care has been taken to ensure the quality of the software. Several unit test-cases have been implemented covering most of the functionality of node and edge addition and deletion. Edge cases concerning degenerate situations like the smallest graph possible that Pytorch Geometric (PyG) can represent, or non admissible actions like re-entering an already existing edge were covered by dedicated Quality Management (QM) Software written in pytest^3^. Furthermore, property-based test-cases [14] were written with the use of the package hypothesis^4^[23]. Those test cases did not explicitly check all the specific changes that are expected to happen in the network structure after an action, but rather the properties and constraints that have to be fulfilled. For example, removing a node makes the number of edges in the network lower or equal to the previous value, makes the number of rows of the array that contains the node features smaller by one, and so on. Correspondingly, removing an edge should keep all information about the nodes alone unchanged; this is an invariant that can be eloquently expressed by the code of a property-based test. Added to that, the selected nodes and edges are not predefined but randomly chosen by the framework. Property-based testing has been proven beneficial for other machine learning problems too [4], [36].

## 3 Results

### 3.1 Application to KIRC dataset

We have verified a potential disease module in the PPI using the Python package GNN-SubNet [29]. The resulting subnetwork or subgraph consists of four proteins, namely MGAT5B, MGAT5, MGAT4B, and MGAT3, connected by five edges (see Figure 7). We have uploaded this module to CLARUS in order to conduct an in-depth investigation of the performance and the distribution of the associated relevance scores. Overall, the module had a high sensitivity of 0.92 and a specificity of 0.42 on an independent test data set (127 patients).

**Figure 7:**
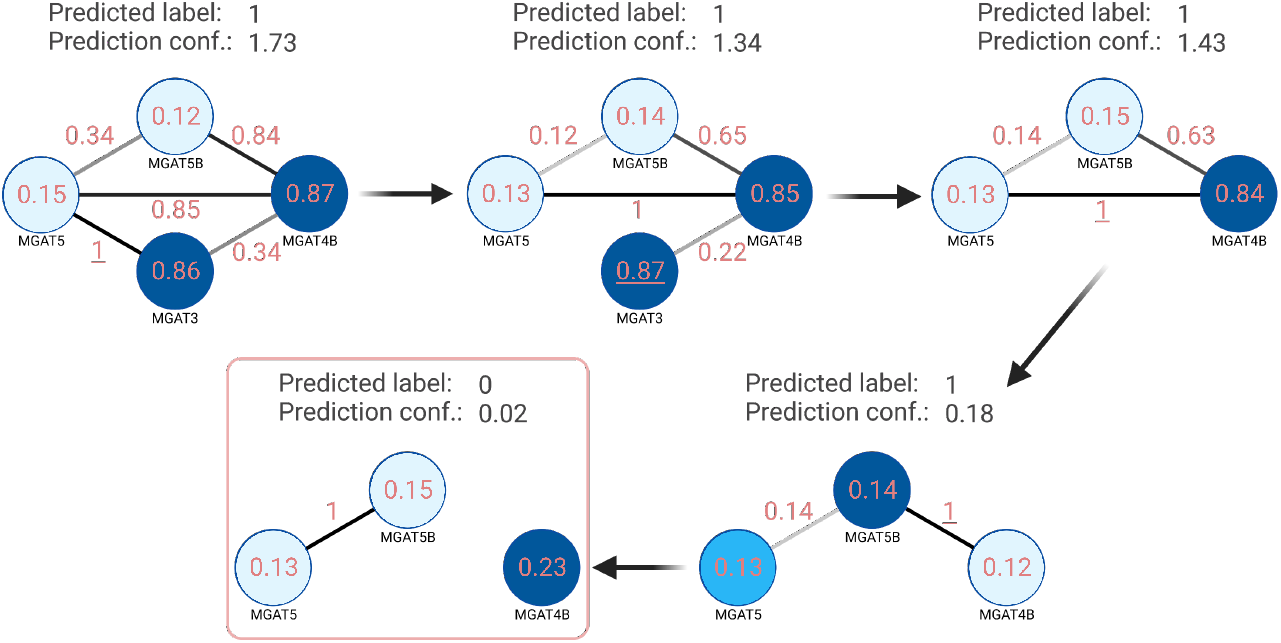
Example manual counterfactual manipulation on the KIRC SubNet dataset of patient graph 62, which had a high predictive confidence of 1.73 in favour of the kidney cancer-specific class. Node relevances are calculated by GNNExplainer [53] and edge relevances by the Integrated Gradients (IG) method [44]. Subsequently, the most relevant edges and nodes are deleted followed each time by a new prediction with the GNN and re-calculation of XAI relevances. This ultimatly results in a counterfactual, a switch in the predicted label from 1 (kidney cancer-specific) to 0 (not kidney cancer-specific), demonstrating the importance of our platform in finding key pathways in the data.

We have selected a patient which was classified with a high predictive confidence of 1.73 (see Table 1) in favour of the kidney cancer-specific class.

**Table 1:** GNN prediction confidence of patient while changing the patient graph.

|  | No changes | Delete edge<br>MGAT3-MGAT5 | Delete node<br>MGAT3 | Delete edge<br>MGAT4B-MGAT5 | Delete edge<br>MGAT4B-MGAT5B |
| --- | --- | --- | --- | --- | --- |
|  | Step 0 | Step 1 | Step 2 | Step 3 | Step 4 |
| Patient Label | 1 | 1 | 1 | 1 | 0 |
| Confidence | 1.73 | 1.34 | 1.43 | 0.18 | 0.02 |

The GNNexplainer assigned the highest relevance score to the node associated with the proteins MGAT3 and MGAT4B (GNNexplainer relevance score of 0.86 and 0.87, see Table 2).

**Table 2:** GNNExplainer importances of nodes while changing the patient graph.

|  | No changes | Delete edge<br>MGAT3-MGAT5 | Delete node<br>MGAT3 | Delete edge<br>MGAT4B-MGAT5 | Delete edge<br>MGAT4B-MGAT5B |
| --- | --- | --- | --- | --- | --- |
| Node | Step 0 | Step 1 | Step 2 | Step 3 | Step 4 |
| MGAT3 | 0.86 | 0.87 | - | - | - |
| MGAT5B | 0.12 | 0.14 | 0.15 | 0.14 | 0.15 |
| MGAT4B | 0.87 | 0.85 | 0.84 | 0.12 | 0.23 |
| MGAT5 | 0.15 | 0.13 | 0.13 | 0.13 | 0.13 |

The edge between MGAT3 and MGAT5 had the highest saliency and IG relevance scores, 1.0 for each XAI method (see Table 3).

**Table 3:** Integrated Gradients (IG) and Saliency (Sal) importances of edges while changing the patient graph.

|  | No changes |  | Delete edge<br>MGAT3-MGAT5 |  | Delete node<br>MGAT3 |  | Delete edge<br>MGAT4B-MGAT5 |  | Delete edge<br>MGAT4B-MGAT5B |  |
| --- | --- | --- | --- | --- | --- | --- | --- | --- | --- | --- |
|  | Step 0 |  | Step 1 |  | Step 2 |  | Step 3 |  | Step 4 |  |
| Edge | IG | Sal | IG | Sal | IG | Sal | IG | Sal | IG | Sal |
| MGAT3-MGAT5 | 1.0 | 1.0 | - | - | - | - | - | - | - | - |
| MGAT3-MGAT4B | 0.34 | 1.0 | 0.22 | 0.55 | - | - | - | - | - | - |
| MGAT4B-MGAT5 | 0.85 | 0.76 | 1.0 | 1.0 | 1.0 | 1.0 | - | - | - | - |
| MGAT4B-MGAT5B | 0.84 | 0.92 | 0.65 | 0.62 | 0.63 | 0.55 | 1.0 | 1.0 | - | - |
| MGAT5-MGAT5B | 0.34 | 0.63 | 0.12 | 0.17 | 0.14 | 0.27 | 0.14 | 0.36 | 1.0 | 1.0 |

Our first manual counterfactual aimed at evaluating the importance of the edge with highest relevance score, MGAT3-MGAT5. The deletion of this edge led to a confidence drop of the predicted class to 1.34. Moreover, a notably higher importance was now assigned to the edge MGAT5-MGAT4B with a saliency and IG relevance score of 1.0 (see Table 3). Interestingly, node importances stayed nearly unchanged for nodes MGAT3 and MGAT4B.

The obtained results may suggest that MGAT3 and MGAT4B are redundant. However, given the remaining topology of the graph we may assume that MGAT4B is more important due to a higher degree than the other nodes. Following this assumption our next manual counterfactual was the deletion of the MGAT3 node. As a result the prediction confidence increased again to 1.43 and the node importance of MGAT4B still barely changed (from 0.85 to 0.84, see Table 2). The importance of the MGAT5-MGAT4B edge stayed the same with a saliency and IG relevance scores of 1.0. The edge between MGAT5 and MGAT4B thus was assumed to be the most relevant part of the analysed module.

According to this assumption and to validate our hypothesis we further deleted the edge between MGAT4B and MGAT5, which resulted in a drastic confidence drop to 0.18 (see Table 1). The importance of MGAT4B decreased to 0.12. The results obtained from this experiment suggest that the information flow through MGAT4B and MGAT5 is crucial for the classification of the analysed patient. MGAT5 as a single marker is not sufficient.

In our last manual counterfactual step we deleted the edge between MGAT5B and MGAT4B, which finally resulted in a counterfactual, where the label switched from 1 (kidney cancer-specific) to 0 (not kidney cancer-specific).

In summary, the conducted experiment demonstrates that human interactions can aid to uncover the most important and relevant parts of a patient graph when guided by recomputed relevance scores after each human counterfactual interaction.

## 4 Discussion

In the Results section we showed how CLARUS can be successfully used by a human-in-the-loop to find the most relevant parts of a patient-specific PPI network through manual counterfactuals. A similar work dealing with explanations of GNNs via targeted topology perturbation is presented in [42]. The “GNNViz” platform belongs to the category of Human-in-the-Loop Machine Learning Systems [15]. It was created to enable non-expert users to manipulate graphs and thereby influence the GNN’s prediction outcome. The benchmark demonstrated that users could uncover label correlations, unfairness, and biases, and a partial comparison with the results of GNNExplainer [53] can be made. In contrast to the “GNNViz” which only considers edge addition and deletion, CLARUS allows and encourages the user to evaluate a variety of manual counterfactuals in the graph such as node addition and deletion. Furthermore, CLARUS is the first tool that allows not only a re-prediction of classes, but also a retraining of the entire GNN after changing the underlying graphs, while other systems such as “GNNViz” solely allow for a recalculation of the prediction result. Other platforms, such as “MetaRelSubNetVis” [1] focus solely on visualisation of patient-specific attributes. It has been used already in the context of XAI to visualise relevance scores within patient specific subnetworks [6]. Nevertheless, the constant jumping between training of AI models and generating XAI relevances to a visualisation platform could potentially be very tedious and time consuming. CLARUS circumvents this by combining visualisations, training of AI methods, and calculation of XAI relevances into one platform resulting in a seamless workflow.

While CLARUS implements a multitude of interactive functionalities, there are other features, which we plan to integrate in future updates of our platform. For instance, the addition and removal of entire node and edge features for all samples. From a usability standpoint this is a valuable feature since some features may be very noisy and thus the user might want to remove that feature completely before retraining. Moreover, the current prototype solely enables the in-depth analysis of a limited set of predefined datasets. In the future, we aim to provide a larger collection of public datasets and the possibility to upload individual datasets of a specific structure along with a user management. This will allow domain-experts to investigate their own datasets within CLARUS. While this is a complex endeavour, involving different implantation and security risks, we have already identified the intricate parts of this feature and plan to integrate it in the next update of CLARUS. Finally, we plan to integrate different types of GNN architectures like the Graph Isomorphism Network (GIN) [51], Jumping Knowledge Network [52], as well as support of a larger variety XAI methods, such as Graph Layer-wise Relevance Propagation (GLRP) [6].

## 5 Conclusion

A lack of trust in AI models in the medical field has prompted the implementation of XAI methods. But to increase trust in AI models we need to move towards causability. Interactive XAI platforms can help bridge the gap between explainability and causability by guiding the human domain expert through the explanations.

We are presenting the first interactive XAI platform prototype, CLARUS, that allows not only the evaluation of specific human counterfactual questions based on user defined alterations of patient-specific PPI networks and a re-prediction of classes, but also a retraining of the entire graph neural network after changing the underlying PPI network structures. Moreover we show the usability of the prototype with the aid of multiple manual counterfactual alterations on a kidney cancer-specific protein-protein interaction network. This example demonstrates the benefit of CLARUS in terms of knowledge gain and causal understanding of the user, for instance on predictions and their explanations.

## 6 Acknowledgements

Dr. Martin Leandro Paleico from the GWDG, the joint data center of Max Planck Society for the Advancement of Science (MPG) and University of Göttingen, assisted our team to transfer and host CLARUS on the GWDG Scientific Compute Cluster. Figures 1 and 7 were created with BioRender.com.

## 8 Abbreviations

AI: Artificial intelligence
CDSS: Clinical decision support system
XAI: Explainable artificial intelligence
ML: Machine learning
PPI: Protein-protein-interaction
GNN: Graph neural network
UI: User interface
KIRC: Kidney Renal Clear Cell Carcinoma
TCGA: The cancer genome atlas
MI: Mutual information
IG: Integrated gradients
PKL: Pickle (file format)

## 9 Availability of data and materials

Project Homepage: http://rshiny.gwdg.de/apps/clarus/ The datasets generated and analysed during the current study are available in python pickle file format (PKL) in the CLARUS platform under the “Select Data”-tab, http://rshiny.gwdg.de/apps/clarus/datasets.zip. A PKL file is a file created by pickle, a python module that enables objects to be serialised to files on disk and de-serialised back into the program at run-time.It contains a byte stream that represents the objects. In this format the datasets can be easily loaded into python and analysed further by the user. Inside the PKL file the data is saved as a list, containing every patient graph. The code for the R frontend is publicly available in a GitHub repository, https://github.com/JacquelineBeinecke/xAI-Shiny-App. The code for the Python backend is publicly available in a GitHub repository, https://github.com/asaranti/GNN_Counterfactuals.

## 11 Authors’ contributions

AH and ACH conceptualised the project. JMB was responsible for the implementation of the frontend in R, as well as, the implementation of the API in Python. AS was responsible for the implementation of the backend in Python. AA implemented the user-management and additionally some API functionality. BP was responsible for the dataset generation and together with AS worked on the application example of the platform. VK implemented a first prototype of the frontend as part of her master thesis. All authors performed review and editing.

## Supplementary

### Screenshots of CLARUS platform

**Figure 8:**
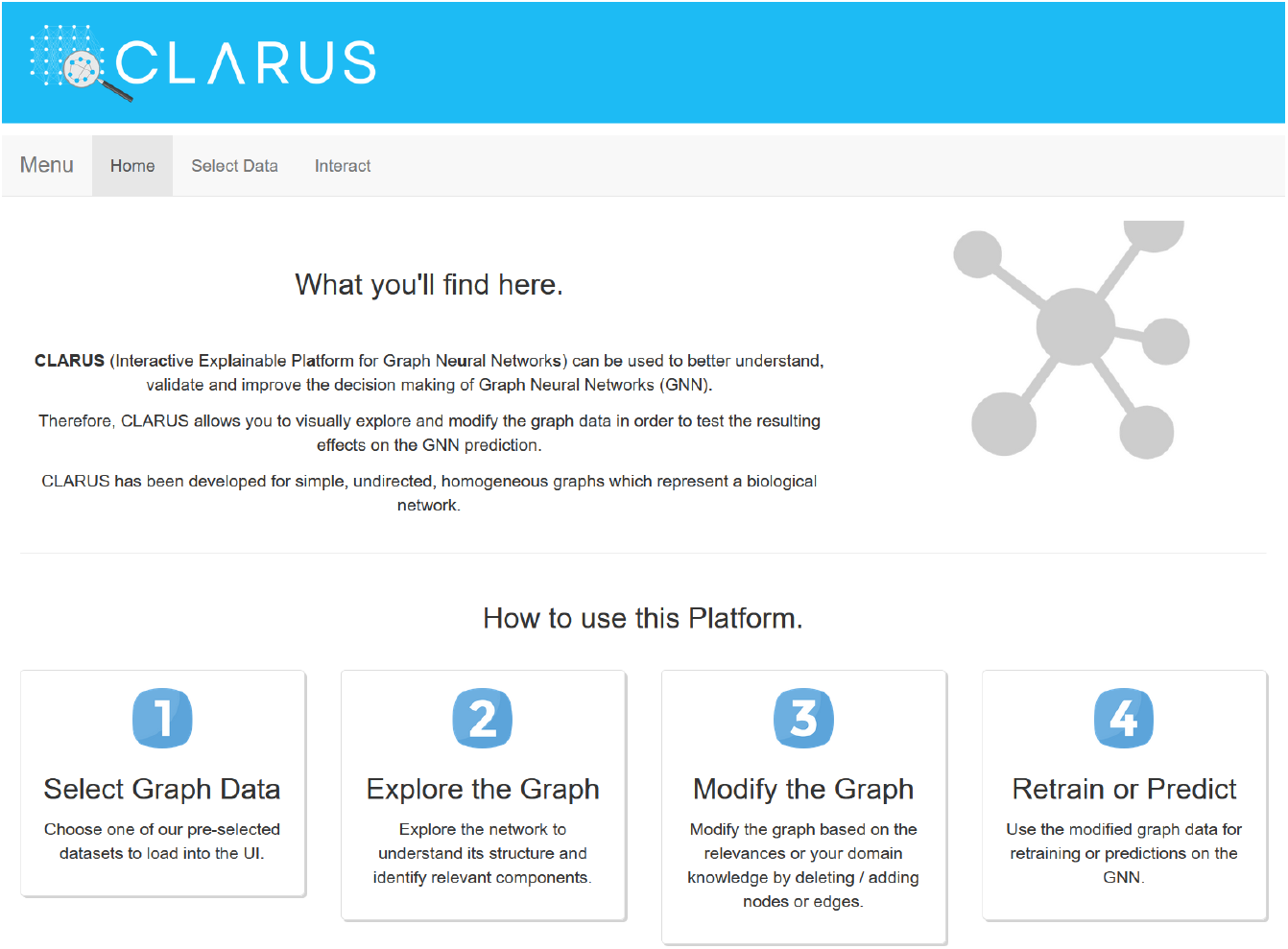
Home screen of the CLARUS platform. This is the first screen a user sees upon opening the platform.

**Figure 9:**
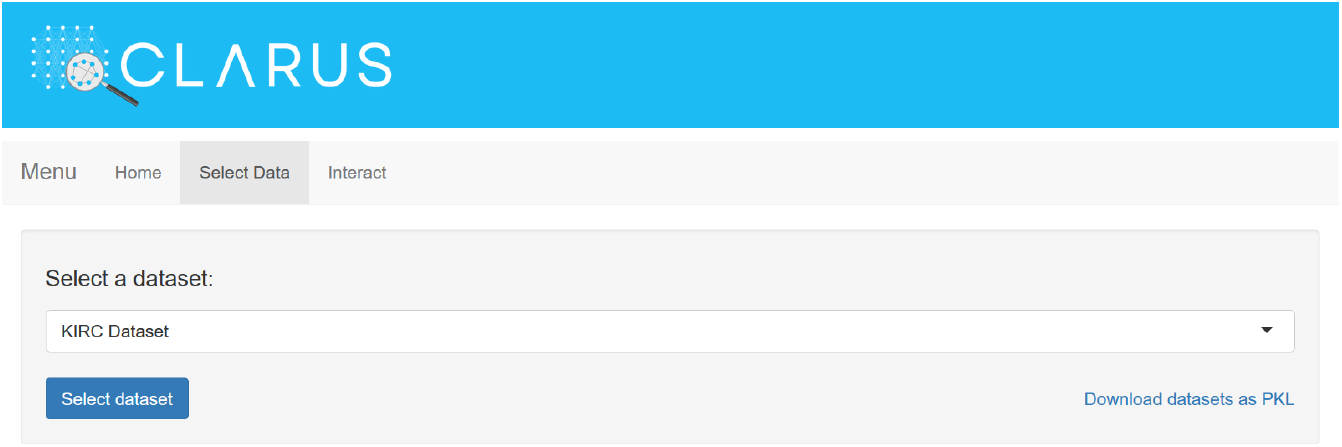
Data selection screen of the CLARUS platform. Here the user can select the dataset they want to analyse and also download the datasets as .pkl files.

**Figure 10:**
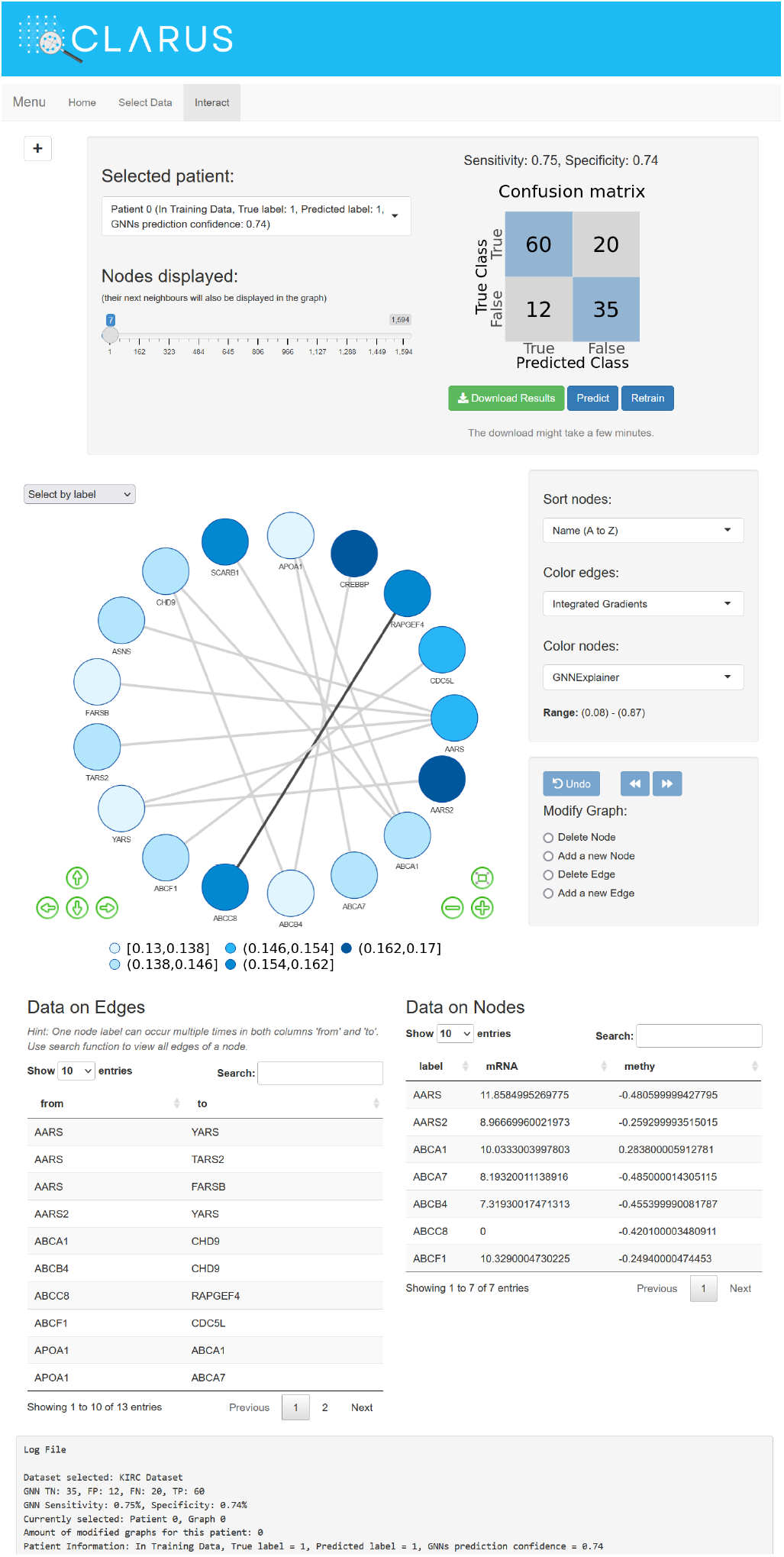
Main window of the CLARUS platform. Here the user is presented all information on the data, alongside XAI attributions and model performance. The user can here manipulate the data, perform a new prediction or retrain the whole model. All manipulations will also be saved verbatim in the log-file, which can also be downloaded.

## Footnotes

1 https://cancergenome.nih.gov/

2 https://captum.ai/

3 https://pytest.org

4 https://hypothesis.readthedocs.io/en/latest/

